# Proteolytic cleavage of the extracellular domain affects signaling of parathyroid hormone receptor 1

**DOI:** 10.1101/2021.12.16.472984

**Authors:** Christoph Klenk, Leif Hommers, Martin J. Lohse

**Author notes:** Department of Biochemistry, University of Zurich, Zurich, Switzerland (Christoph Klenk). Corresponding author: Christoph Klenk.

## Abstract

Parathyroid hormone 1 receptor (PTH1R) is a member of the class B family of G protein-coupled receptors, which are characterized by a large extracellular domain required for ligand binding. We have previously shown that the extracellular domain of PTH1R is subject to metalloproteinase cleavage *in vivo* that is regulated by ligand-induced receptor trafficking and leads to impaired stability of PTH1R. In this work, we localize the cleavage site in the first loop of the extracellular domain using amino-terminal protein sequencing of purified receptor and by mutagenesis studies. We further show, that a receptor mutant not susceptible to proteolytic cleavage exhibits reduced signaling to Gs and increased activation of Gq compared to wild-type PTH1R. These findings indicate that the extracellular domain modulates PTH1R signaling specificity, and that its cleavage affects receptor signaling.

## Introduction

Parathyroid hormone 1 receptor (PTH1R) is a key regulator of blood calcium levels and bone metabolism in response to parathyroid hormone (PTH). Moreover, activation of PTH1R by parathyroid-related hormone peptide (PTHrP) has been implicated in fetal development and in malignancy-associated hypercalcemia (1). PTH1R is a member of the class B family of G protein-coupled receptors (GPCRs) which are characterized by a large N-terminal extracellular domain (ECD; ~100 to 180 residues) that is critically involved in ligand binding (2,3). Similar to other class B GPCRs, the ECD of PTH1R consists of two pairs of antiparallel β-strands flanked by a long and a short α-helical segment at the N- and C-terminal end, respectively. The overall conformation is constrained by three conserved disulfide bonds which are required for proper folding and for ligand binding (4–7). The ECD is oriented in an upright position above the membrane surface with residues 15-34 of PTH binding into a groove formed by the ECD and the N-terminal part of the ligand protrudes as a continuous α-helix into the transmembrane domain of PTH1R (Fig. 1) (7). In line with the receptor structure, a two-step activation model has been proposed, where first the C-terminal part of PTH binds to the extracellular domain, and then the N-terminal part of PTH interacts with the receptor core, thereby leading to receptor activation (8,9). PTH1R couples to multiple heterotrimeric G protein subtypes and can activate several signaling pathways concomitantly. Predominantly, adenylyl cyclases are stimulated by activation of G_s_ as well as phospholipase Cβ by G_q_ (10–12). Moreover, activation of G_i/o_ resulting in inhibition of adenylyl cyclase and activation of G_12/13_ leading to phospholipase D and RhoA activation, as well as activation of mitogen-activated protein kinases through G protein-dependent and -independent mechanisms have been reported (13–18). PTH1R activation can have anabolic and catabolic effects on bone. While continuous administration of PTH enhances osteoclastogenesis leading to bone resorption and calcium liberation, intermittent administration of PTH results in bone formation through enhancing osteoblast differentiation and survival, which is used as a treatment option for severe osteoporosis (19,20). Although the exact molecular mechanisms are not clear yet, differential activation of signaling pathways seems to play an important role in these opposing effects upon PTH1R activation. While G_s_-signaling is the predominant pathway for promoting PTH-induced bone formation, G_q_-activation seems to have little or no effect on osteogenesis (21–23). In addition, β-arrestin recruitment was shown to be essential for selectively promoting bone formation upon treatment of mice with recombinant PTH(1–34) (24). Moreover, many of these effects appear to be regulated in a tissue- and cell-type specific manner (25).

**Figure 1.**
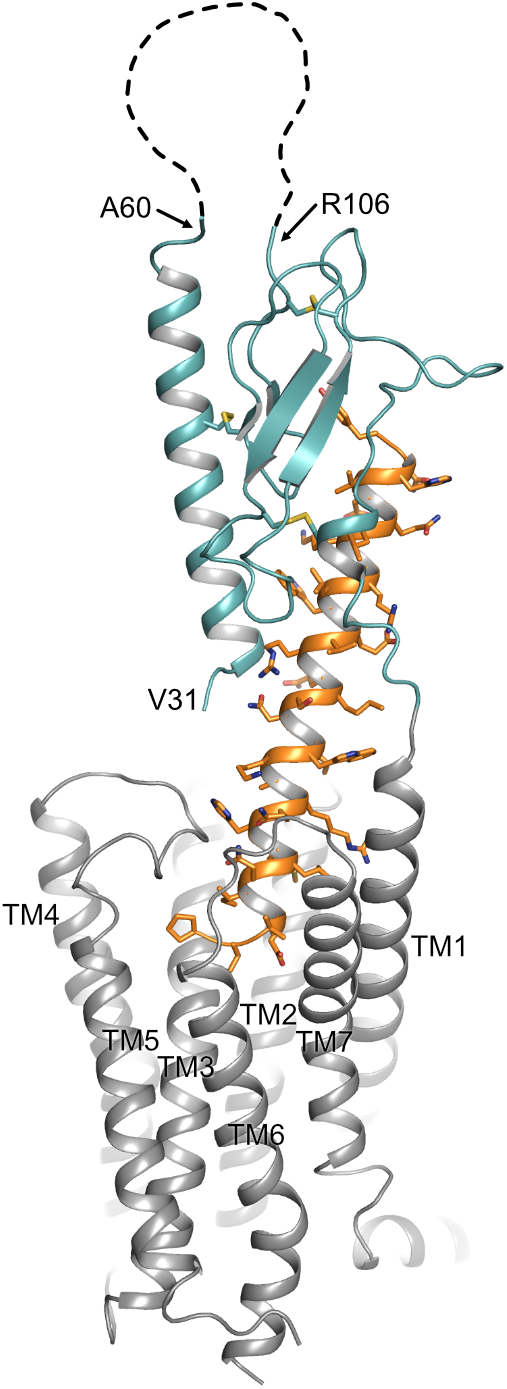
Topology of PTH1R. Structure of the human PTH1R (transmembrane domain, grey; ECD, teal) in complex with a PTH analog (orange) (PDB ID: 6FJ3). Unstructured residues 61-105 of ECD loop 1 are depicted as a dashed line. The receptor N-terminal residue (V^31^, as resolved in the crystal structure), residues embracing ECD loop1, and transmembrane helices (TM1-7) are indicated.

We have previously shown that the ECD of PTH1R can undergo proteolytic cleavage by an extracellular metalloproteinase resulting in reduced stability and degradation of the receptor. We also demonstrated that N-terminal ECD cleavage occurred only at the cell surface, and that internalization of the receptor resulting from continuous activation by agonists prevented cleavage and thereby stabilized the receptor. Furthermore, our experiments suggested that the cleavage site is located within the first 90 residues of the receptor, however the exact position was not fully resolved (26). In the present study, we have localized the cleavage site in the unstructured loop of exon E2 within the PTH1R ECD. Moreover, we demonstrate that ECD cleavage results in an altered ligand efficacy of PTH changing the G protein-coupling of PTH1R from G_q_ to G_s_.

## Materials and Methods

### Materials

Lipofectamine 2000 was purchased from Thermo Fisher Scientific. [Nle^8,18^,Tyr^34^]PTH (1–34), a chemically more stable variant of native PTH, was purchased from Bachem and is referred to as PTH(1–34). Generation of a polyclonal rabbit anti-PTH1R (1781) antiserum was described previously (27). Anti-rabbit peroxidase-conjugated secondary antibodies were obtained from Dianova. Cy2-conjugated anti-rabbit antibody was from Jackson Immuno Research Lab. All cell culture media were obtained from PAN Biotech. All other reagents were of analytical grade from Sigma-Aldrich or Applichem.

### cDNA constructs

A Strep-Tag II (WSHPQFEK) was fused to the C-terminal end of human PTH1R (26) by PCR. Alanine mutations were introduced into the extracellular domain of PTH1R by overlap extension PCR. In total, 6 constructs with alanine blocks from Leu^56^-Met^63^, Glu^64^-Ser^71^, Ala^72^-Arg^79^, Lys^80^-Leu^87^, Tyr^88^-Lys^95^, and Glu^96^-Tyr^103^ were generated. All constructs were subcloned into pcDNA5/FRT vector (Thermo Fisher Scientific) using the restriction sites EcoRI and ApaI and verified by sequencing.

### Cell culture and transfection

Flp-In CHO cells (Thermo Fisher Scientific) were maintained in 1:1 Dulbecco’s modified Eagle’s medium/ Ham’s F12 medium containing 10 % (v/v) fetal calf serum, 100 U/ml penicillin, 100 μg/ml streptomycin and 100 μg/ml Zeocin (Thermo Fisher Scientific). Cells were maintained at 37 °C in a humidified atmosphere of 5% CO_2_, 95% air. To generate stable cell lines, cells were transfected with pcDNA5/FRT-PTH1R plasmids using Lipofectamine 2000 according to the manufacturer’s instructions. 48 h after transfection, cells were selected in culture medium where Zeocin was replaced by 600 μg/ml hygromycin B for approximately two weeks. Clonal cell lines were derived from limited dilution series and screened for expression of PTH1R by Western blot and immunocytochemistry.

### SDS-PAGE and Western blotting

Cells were lysed in SDS-loading buffer [50 mM Tris (pH 6.8), 2% (v/v) SDS, 10% glycerol, 5 mg/ml bromophenol blue] for 20 min on ice, briefly sonified and incubated at 45 °C for 20 min. For reducing conditions, 4% (v/v) β-mercaptoethanol was added to the lysis buffer. Lysates were cleared by centrifugation and run on 10% SDS-polyacrylamide gels in a Mini-PROTEAN 3 cell apparatus (Biorad). Proteins were electroblotted onto Immobilon P membranes (Millipore) using a Bio-Rad Mini trans-blot cell apparatus at 100 V for 60 min at 4 °C. The blots were probed with anti-PTH1R antibodies (1:4,000), followed by horseradish peroxidase-conjugated goat anti-rabbit (1:10,000) and detection on Super RX X-ray film (Fujifilm) using ECL Plus reagent (GE Healthcare).

### Purification of PTH1R

Membranes from Flp-In CHO cells stably expressing PTH1R-Strep2 were prepared as described before (28). Membranes were solubilized in solubilization buffer [50 mM Tris-HCl (pH 7.4), 140 mM NaCl, 0.5% (w/v) n-dodecyl β-D-maltoside, 10 μg/ml soybean trypsin inhibitor, 30 μg/ml benzamidine, 5 μg/ml leupeptin, 100 μM PMSF] for 2 h, and insoluble material was removed by centrifugation for 1 h at 100,000 × g. Solubilized PTH1R-Strep2 was incubated with 1 ml Strep-Tactin sepharose (IBA GmbH) for 12 h under constant agitation and washed with 10 column volumes of wash buffer [100 mM Tris-HCl (pH 8.0), 1 mM EDTA, 150 mM NaCl, 0.1% (w/v) n-dodecyl β-D-maltoside]. Bound receptor was eluted with 1-2 column volumes of the same buffer supplemented with 2.5 mM desthiobiotin and concentrated with a Microcon centrifugal filter device (10,000 MWCO, Millipore).

### Amino-terminal sequencing of cleaved receptor fragments

Fifty μg of purified receptor fragments were blotted onto PVDF membranes and stained with Coomassie Blue. The fragments were excised and subjected to automated Edman degradation (Wita GmbH).

### Immunocytochemistry and confocal imaging

CHO cells stably expressing PTHR variants were grown on coverslips overnight. Cells were then exposed (or not) to 100 nM PTH(1–34) for 30 min as indicated. Cells were fixed with 4% paraformaldehyde and 0.2% picric acid in 0.1 M phosphate buffer (pH 6.9) for 30 min at room temperature and washed five times in PBS. For permeabilization cells were incubated for 5 min in methanol. After 10 min of preincubation in PBS containing 0.35% (w/v) BSA, cells were incubated with anti-PTH1R antibody at a dilution of 1:2,000 in PBS containing 0.35% (w/v) BSA for 1 h at 37 °C. Bound primary antibody was detected with Cy2-labeled goat anti-rabbit IgG (1:400). Specimens were examined using a Leica SP2 laser scanning confocal microscope.

### Functional receptor assays

Signaling assays were measured in Flp-In CHO cells stably expressing the indicated PTH1R variants. cAMP was measured using a RIA kit (Beckman Coulter), and inositol phosphates were separated by chromatographic separation of myo-[2-^3^H]inositol phosphates as described previously (29). Pharmacological data were analyzed in Prism v6.0 (GraphPad Software). A three-parameter logistic equation was fit to the data to obtain concentration-response curves and E_max_ values. Statistical differences were analyzed using unpaired t-tests.

## Results

Our previous findings suggested that the protease cleavage site is most likely located between Cys^48^ and Cys^108^ (26), a region including exon E2 of PTH1R which is unique among all other class B GPCRs and which forms a disordered loop in the crystal structures of the isolated PTH1R ECD (4) and full length PTH1R (7) (Fig. 1). We therefore created a series of mutants where stretches of 8 residues in this region were mutated to Ala to delineate the protease cleavage site (Fig. 2A). Each of the six resulting PTH1R variants was stably expressed in Flp-In CHO cells, and expression and membrane targeting of the receptor were analyzed by immunofluorescence (Fig. 2B) using an antibody detecting the C-terminal part of PTH1R (26,27). All mutants exhibited a distinct membrane staining which was comparable to that of wild-type receptor. Upon stimulation with 100 nM PTH(1–34) for 30 min, a sequestration of receptor from the cell surface into endocytic vesicles was observed suggesting that each receptor variant activated intracellular signaling pathways leading to receptor internalization.

**Figure 2.**
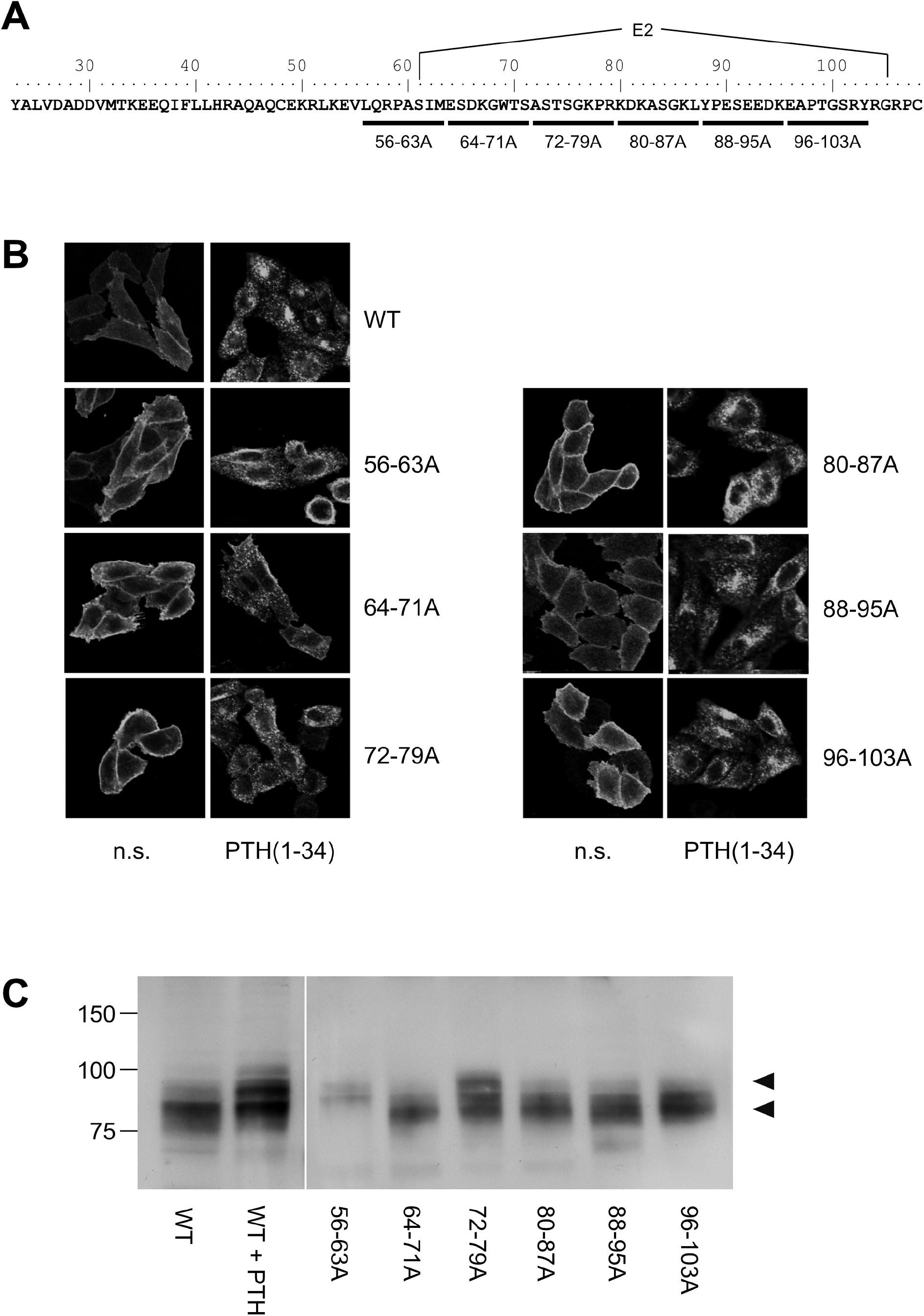
Mapping of the protease cleavage site of PTH1R ECD by alanine scan. **(A)** Amino acid sequence of the extracellular domain of the human PTH1R. Stretches of 8 amino acids that were replaced by alanine residues are indicated by horizontal bars. Exon E2 is marked above the sequence. **(B)** CHO cells stably expressing PTHR variants were treated with 100 nM PTH(1–34) for 30 min or left untreated. Subsequently, cells were fixed, permeabilized, and stained with rabbit anti-PTH1R antibody followed by Cy2-labeled anti-rabbit antibody. PTHR was visualized by confocal microscopy. **(C)** CHO cells stably expressing PTHR variants were lysed, and PTH1R was monitored by reducing SDS-PAGE and Western blotting. Cells were treated with 100 nM PTH(1–34) for 12 h prior to cell lysis where indicated. Arrowheads depict the cleaved (MW ~80 kDa) and the uncleaved (MW ~90 kDa) PTH1R band.

To test whether any of the mutations had an effect on protease cleavage, the migration patterns of PTH1R variants were analyzed by reducing SDS-PAGE and Western blotting. As demonstrated previously, the ECD of PTH1R residing at the cell surface is cleaved by extracellular metalloproteinases. The resulting N-terminal fragment of the ECD (~10 kDa) remains tethered to the receptor core by a single disulfide bond under native conditions but is lost under reducing conditions leading to an apparent molecular weight of the receptor of ~80 kDa (26). In contrast, sustained receptor activation by PTH(1–34) resulting in continuous receptor internalization rendered the receptor inaccessible for extracellular proteases and thus protected the full-length receptor with a molecular weight of ~90 kDa (Fig 2C, left panel; c.f. (26)). Similar to wild-type PTH1R, mutants where residues 64-71, 80-87, 88-95 or 96-103 had been replaced by alanines migrated at ~80 kDa in reducing SDS-PAGE, indicating that protease cleavage was not prevented by the respective mutations. In contrast, PTH1R^56-63A^ exclusively migrated at ~90 kDa similar to wild-type receptor where cleavage had been prevented by stimulation with PTH(1–34). For PTH1R^72-79A^ two bands were detected which co-migrated with the cleaved and non-cleaved receptor species. Thus, mutating the region between Leu^56^ and Met^63^ to alanine fully inhibited proteolytic cleavage of the PTH1R ECD, indicating that the main cleavage site is located in this region. PTH1R^72-79A^ exhibited incomplete inhibition of cleavage, suggesting another, less susceptible cleavage site or an incomplete masking of the cleavage site around residues 56-63.

To corroborate these findings, we determined the amino acid sequence of the N-terminus of the cleaved 80 kDa fragment of PTH1R. PTH1R was purified from stably expressing Flp-In CHO cells via a C-terminal Strep2 tag. 50 μg of purified protein were blotted onto PVDF membrane and stained with Coomassie blue. A band corresponding to the cleaved PTH1R fragment was excised and analyzed by Edman degradation (Fig. 3A). Three different N-termini were identified, located at positions Ser^65^, Ser^73^ and Lys^80^ in the ECD of PTH1R (Fig. 3B). Finally, the complete ECD of PTH1R (residues 23-177) was subjected to a computational cleavage site search using positional weight matrices (PWM) for 11 matrix metalloproteinases (MMPs) (30). This procedure revealed a total of 19 putative cleavage sites located between residues 30 to 173. However, only cleavage site Ser^61^↓Ile^62^ was common to all 11 MMPs and exhibited the highest PWM scores among all other predicted cleavage sites (Table 1). Taken together, these findings support the results of the alanine scan, suggesting that the primary cleavage occurs at Ser^61^↓Ile^62^ of PTH1R.

**Figure 3.**
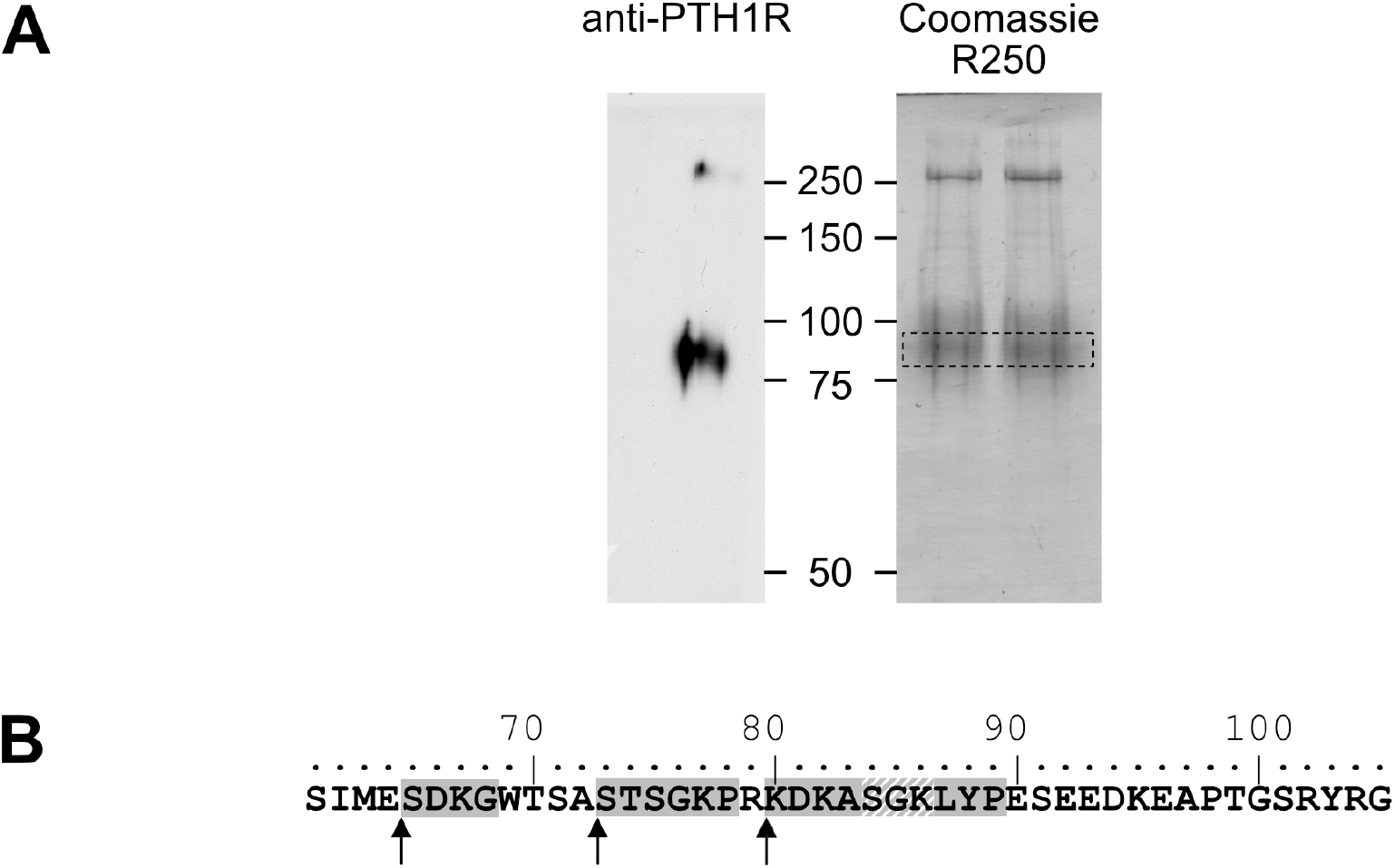
Mapping of the protease cleavage site of PTH1R ECD by N-terminal sequencing. **(A)** Human PTH1R with C-terminal Strep2-tag was purified from stably expressing CHO cells by two-step affinity purification. A fraction of the purified receptor was subjected to Western blot and probed with anti-PTH1R antibodies (left panel). The remaining purified receptor protein (~50 μg) was transferred on PVDF membranes and stained with Coomassie blue R250. The band corresponding to PTH1R was cut out and subjected to microsequencing (right panel, dashed box). **(B)** Sequence of exon E2 (amino acids 61-105). Sequences obtained from microsequencing are shaded gray. The position of the N-terminal amino acid is marked by an arrow. Residues 84-86 (gray diagonal stripes) were not resolved in the Edman degradation.

**Table 1.** Computational cleavage site prediction of PTH1R ECD. The sequence of the mature PTH1R ECD (amino acids 23-177) was analyzed for MMP cleavage sites with CleavPredict (30) using position weight matrices for 11 MMPs (MMP-2, MMP-3, MMP-8, MMP-9, MMP-13, MMP-14, MMP-15, MMP-16, MMP-17, MMP-24, MMP-25). For each cleavage site, the residue number of P1, the sequence corresponding to P5-P5’ (numbering according to Schechter and Berger (62)) and the position weight matrix score (PWM score) for each MMP subtype are given.

| P1<br>position | residues<br>(P5-P5') | MMP |  |  |  |  |  |  |  |  |  |  |
| --- | --- | --- | --- | --- | --- | --- | --- | --- | --- | --- | --- | --- |
|  |  | 2 | 3 | 8 | 9 | 10 | 14 | 15 | 16 | 17 | 24 | 25 |
| 30 | VDADD↓VMTKE |  |  |  |  |  |  |  |  |  |  | 1.32 |
| 37 | TKEEQ↓IFLLH |  |  |  |  |  |  |  |  |  |  | 0.19 |
| 40 | EQIFL↓LHRAQ |  | 3.46 |  |  | 2.19 |  |  |  | 3.43 |  | 1.50 |
| 46 | HRAQA↓QCEKR |  |  |  |  |  |  |  |  | 2.37 |  | 1.71 |
| 51 | QCEKR↓LKEVL |  |  |  |  |  |  |  |  |  |  |  |
| 61 | QRPAS↓IMESD | 6.75 | 8.76 | 6.13 | 5.72 | 8.26 | 7.60 | 5.37 | 7.00 | 3.42 | 7.88 | 5.82 |
| 80 | GKPRK↓DKASG | 1.36 |  |  |  |  |  |  |  |  |  |  |
| 86 | KASGK↓LYPES |  |  | 1.63 |  |  |  | 1.31 |  | 2.44 |  | 4.71 |
| 100 | EAPTG↓SRYRG |  |  |  | 2.52 |  |  |  |  |  |  |  |
| 102 | PTGSR↓YRGRP |  |  | 1.90 |  |  |  |  |  |  |  |  |
| 114 | PEWDH↓ILCWP |  |  |  |  |  |  | 2.62 |  |  |  |  |
| 125 | GAPGE↓VVAVP | 3.56 | 3.14 | 2.03 |  |  |  |  |  |  |  | 0.21 |
| 134 | PCPDY↓IYDFN |  |  | 2.48 |  | 1.64 |  | 5.02 |  |  |  | 4.48 |
| 137 | DYIYD↓FNHKG |  |  | 2.06 |  |  |  |  |  |  |  |  |
| 144 | HKGHA↓YRRCD |  |  | 2.67 |  |  |  |  |  |  | 0.81 |  |
| 155 | NGSWE↓LVPGH |  |  | 5.18 |  |  |  |  |  | 2.47 |  | 4.00 |
| 160 | LVPGH↓NRTWA | 2.05 |  |  |  |  | 1.74 | 2.03 | 1.35 |  |  |  |
| 166 | RTWAN↓YSECV |  |  |  |  |  | 1.97 |  | 2.16 |  |  |  |
| 173 | ECVKF↓LTNET |  |  | 3.15 | 6.44 |  | 1.30 |  | 2.98 | 3.50 |  | 4.39 |

To test whether ECD cleavage affected receptor function, we assessed activation of the two canonical signaling pathways of PTH1R G_s_ (cyclic AMP, cAMP) and G_q_ (inositol phosphates, IP) by wild-type PTH1R, by the fully cleavage-deficient mutant PTH1R^56-63A^, and by the partially cleaved mutant PTH1R^72-79A^. All measurements were performed in stably expressing CHO cells that had been matched for equal receptor expression levels. Compared to wild-type PTH1R, maximal PTH-induced generation of cAMP was reduced by 37% for PTH1R^56-63A^, whereas no change was observed for PTH1R^72-76A^ (Fig. 4A, Table 2). In contrast, PTH-induced generation of [^3^H]IP was increased by 35% for PTH1R^56-63A^, whereas no change was observed for PTH1R^72-79A^ (Fig. 4B, Table 2). In summary, these findings suggest, that full cleavage of the ECD of PTH1R leads to decreased efficacy of PTH(1–34) in G_q_ signaling and increased efficacy in Gs signaling. PTH1R^72-79A^ did not differ from wild-type PTH1R, which may be explained by the fact that the majority of PTH1R^72-79A^ was still proteolytically processed (Fig. 3B). Thus, cleavage appears to directly modulate the signaling bias of PTH1R.

**Figure 4.**
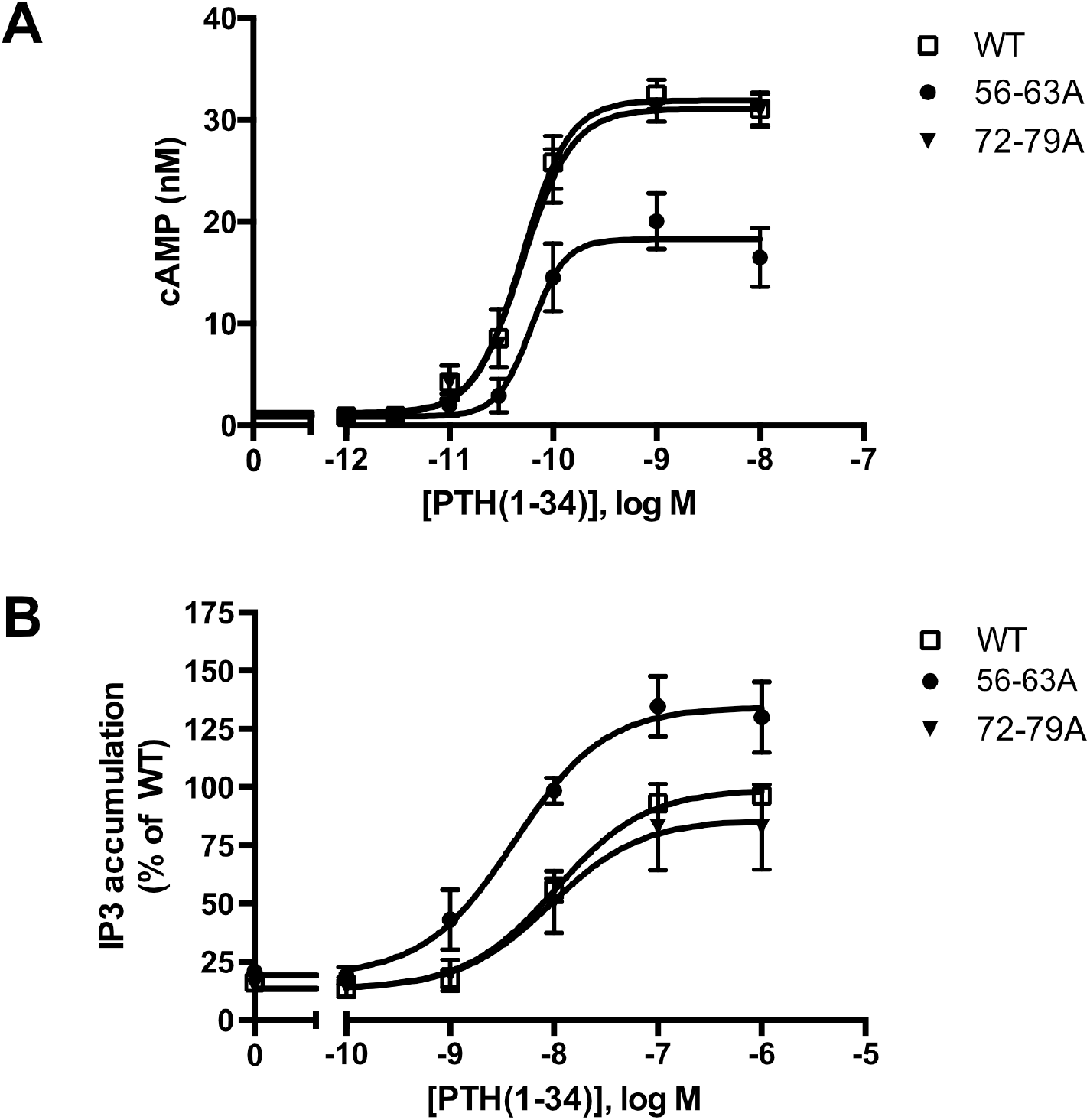
Protease cleavage changes the signaling specificity of PTH1R from G_q_ and G_s_. **(A)** Flp-In CHO cells stably expressing PTH1R, PTH1R^56-63A^ or PTH1R^72-79A^ were stimulated for 20 min with PTH(1–34) at the indicated concentrations, and cAMP levels were quantified with a radioimmunoassay. The means ± S.E.M. of five independent experiments are shown. **(B)** Flp-In CHO cells stably expressing PTH1R, PTH1R^56-63A^ or PTH1R^72-79A^ were incubated with [*myo*-2-^3^H(N)]inositol and 0.2% fetal calf serum for 16 h. Cells were stimulated for 60 min with the indicated concentrations of PTH, and [^3^H]IP_3_ levels were quantified in a scintillation counter after chromatographic separation. Data represent the means ± S.E.M. of five individual experiments.

**Table 2.** Effects of ECD cleavage on cAMP generation and IP accumulation. PTH-induced E_max_ values for cAMP and IP generation of PTH1R^56-63A^ or PTH1R^72-79A^. were compared against that of PTH1R using unpaired t tests. Data summarize results of 3-5 independent experiments.

|  | Difference vs. PTH1R, cAMP (%) |  |  | Difference vs. PTH1R, [<sup>3</sup>H] IP (%) |  |  |
| --- | --- | --- | --- | --- | --- | --- |
|  | Mean ± SEM | 95% C.I. | *p* | Mean ± SEM | 95% C.I. | *p* |
| PTH1R<sup>56-63A</sup> | -37.3 ± 5.9 | -52.7 to -21.9 | 0.0015 | 35.3 ± 10.7 | 10.6 to 60.1 | 0.011 |
| PTH1R<sup>72-79A</sup> | -1.9 ± 1.3 | -5.3 to 1.5 | 0.21 | -5.6 ± 19.4 | -48.8 to 37.6 | 0.78 |

C.I., confidence interval

## Discussion

Previously, we have reported that the ECD of PTH1R is subject to cleavage by metalloproteinases. PTH1R cleavage is a constitutive phenomenon and is inhibited by receptor activation (26). In the present study we aimed to characterize the role of proteolytic processing of the extracellular domain, and we provide evidence for the exact location of the cleavage site as well as for a modulation of signaling properties upon cleavage. N-terminal sequencing of the 80 kDa receptor core (remaining after shedding the cleaved N-terminal fragment by disulfide hydrolysis) revealed three nearby cleavage sites (Glu^64^↓Ser^65^, Ala^72^↓Ser^73^ and Arg^79^↓Lys^80^). However, computational analysis using cleavage patterns of 11 MMPs suggested a putative cleavage site at Ser^61^↓Ile^62^ which was located 3 amino acids upstream of the first free N-terminus identified by microsequencing. A systematic alanine scan within this region of the ECD showed, that only PTH1R^56-63A^ was completely resistant to proteolysis, while mutation of residues 64-71 to alanine did not prevent proteolysis, further supporting the proximal site at residue 61. Only a fraction of PTH1R^72-79A^ remained intact whereas the majority of receptor was found as the cleaved 80 kDa form. This may suggest, that the alanine mutations at residues 72-79 mask the cleavage site around residues 56-63 to some extent or may hamper protease interaction resulting in incomplete protease cleavage. Considering the results from computational and biochemical analyses, we propose that Ser^61^↓Ile^62^ is the most likely primary cleavage site. The free N-termini observed in Edman degradation at Ser^65^, Ser^73^ and Lys^80^ may be the result of limited exopeptidase action following endopeptidase cleavage. Ser^61^ is located within the first residues of a large loop connecting the top layer formed by α1-helix with the first β-strand of the middle layer of the α-β-β-α fold of PTH1R ECD. Notably, residues 61-104 were not resolved in any structure of PTH1R-ECD or full length PTH1R suggesting high flexibility in this region (4,7,31). Considering the orientation of the ECD in the full-length structures, Ser^61^ would be located at the distal part of the receptor facing away from the membrane and, thus, may be well accessible for extracellular proteases (Fig 1).

Processing by MMPs and other metalloproteinases has been described previously for a limited number of other GPCRs, e.g. for β_1_-adrenergic receptor (32), endothelin B receptor (33,34), thyrotropin receptor (35,36), protease-activated receptor 1 (PAR-1) (37,38), GPR124 (39) and more recently for GPR37 (40,41). Apart from PAR-1 and the adhesion family receptor GPR124, where protease cleavage unmasks the endogenous ligand resulting in receptor activation, a functional consequence of protease cleavage has not been explicitly reported. To our surprise, proteolytic cleavage of the PTH1R ECD directly affected receptor signaling. In contrast to wild-type receptor, the cleavage-deficient mutant PTH1R^56-63A^ exhibited reduced cAMP and increased IP responses to PTH stimulation. Protease cleavage thus enhanced coupling efficacy of the receptor to the G_s_ pathway, while reducing G_q_-coupling at the same time, resulting in a signaling bias. Biased signaling is defined as ligands giving different degrees of activation in separate signaling pathways of the same receptor. Besides binding of ligands to allosteric sites on the receptor that stabilize distinct active receptor conformations, interaction of a receptor with intracellular adaptor proteins and subcellular receptor sequestration have been reported to affect signaling bias (42–44). For PTH1R, several ligands and intracellular adaptors which direct signaling specificity to G_s_, G_q_ or G prote-inindependent pathways have been described (17,22,24,45–48). All of these PTH/PTHrP derivatives carry modifications at the N-terminal part, which directly interacts with the transmembrane domain of the receptor, suggesting that signaling specificity is mediated by direct conformational stabilization of the receptor core. Our findings now indicate that the ECD, which accommodates the C-terminal part of PTH and which is commonly believed to only serve as an “affinity trap” for the ligand, can also affect signaling specificity of the receptor.

There is growing evidence, that extracellular regions of GPCRs play important roles in fine-tuning receptor activity and signaling selectivity. Apart from PARs and adhesion receptors, where the buried ligand is proteolytically released from the receptor’s N-terminus, extracellular loops play an important role in modulating the function of several class A and class B receptors (49). More importantly, calcium-mediated interaction of extracellular loop 1 and PTH has been shown to modulate PTH1R activity (50–52). Recent studies on glucagon receptor suggest that the ECD itself may act as an allosteric inhibitor by interaction of α1-helix of the ECD with extracellular loop 3 of the receptor core (53). Moreover, recent cryo-EM structures of active-state class B GPCRs including PTH1R reveal a high degree of conformational flexibility of the ECD (31,54,55), and it has been proposed that the dynamic motion of the ECD may contribute to biased agonism of class B GPCR ligands (56,57). In line with that, an antibody primarily binding to α1-helix of the ECD has been shown to modulate β-arrestin signaling of PTH1R (58) suggesting that perturbation of ECD orientation or conformation may alter receptor signaling. Proteolytic cleavage at Ser^61^ is expected to result in increased conformational flexibility of α1-helix of PTH1R ECD as the helix remains tethered to the receptor only through a disulfide bond between Cys^48^ and Cys^117^(26). As a consequence, especially the N-terminal part of α1-helix may gain additional flexibility (Fig. 1). Notably, within this region residues 32-41 make important contacts to PTH including the flexible central region of the peptide (Fig. 5) (4,7). This region was shown to be critical for initiating the two-step binding mechanism of PTH (59). Thus, it may well be conceivable that alterations in the flexibility and orientation of α1-helix of PTH1R ECD can allosterically affect receptor signaling. Whether these effects are mediated by an altered interaction of the ECD with the transmembrane core, by a rearrangement of the ligand in the binding pocket, or involve interaction of additional proteins such as RAMPs (receptor activity-modifying proteins) with PTH1R (60,61) needs to be studied.

**Figure 5.**
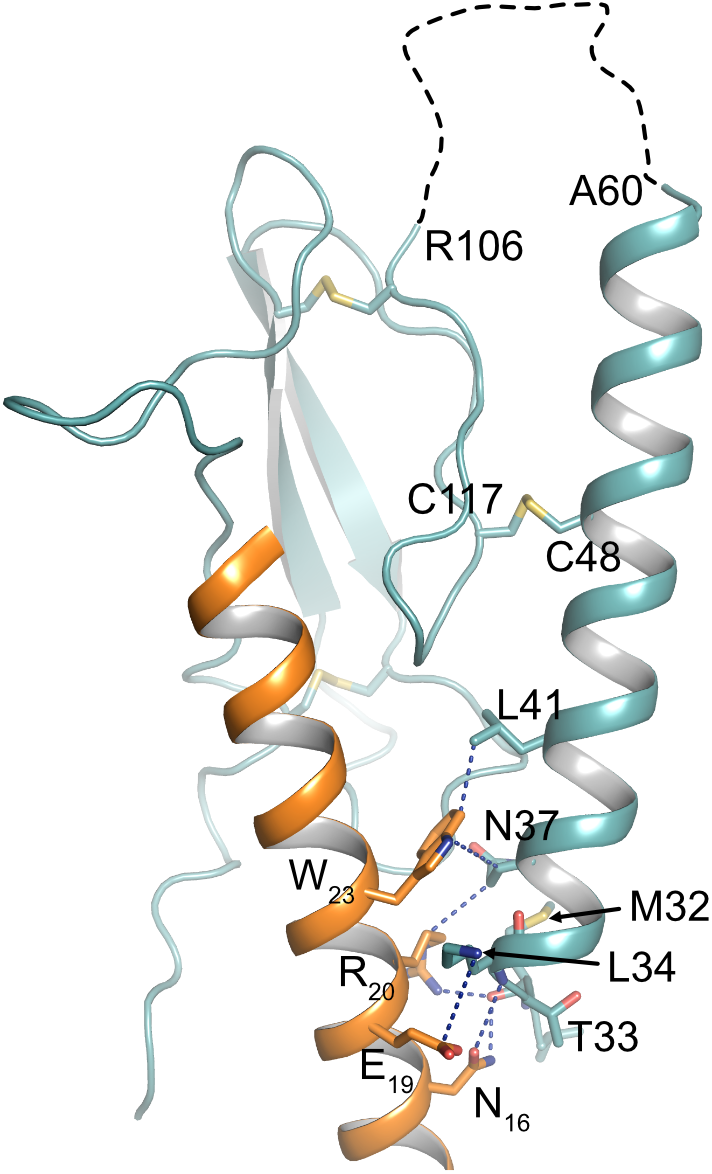
Interface between PTH and α1-helix of PTH1R ECD. Crystal structure of the PTH1R ECD (teal) in complex with PTH (orange) (PDB ID: 6FJ3). Residues forming the interface are shown as stick, and contacts are indicated as blue dashed lines. The unstructured loop 1 of the ECD is shown as black dashed line.

In summary, we have mapped the cleavage site within the ECD of PTH1R and demonstrate for the first time, to our knowledge, that protease cleavage of the ECD of a GPCR modulates G protein signaling specificity.

## Author Contributions

CK and MJL conceived the study; CK designed experiments; CK and LH performed experiments; CK, LH and MJL analyzed data; CK wrote the manuscript, all authors provided edits and comments.

## Acknowledgments

We thank Michaela Hoffmann, Annette Hannawacker, Alexandra Bohl and Monika Frank for technical assistance. Moreover, we thank Dr. Thomas Pohl (Wita GmbH, Berlin) for support with N-terminal microsequencing.

## References

1. Jüppner H, Abou-Samra AB, Uneno S, Gu WX, Potts JT, Segre GV. The parathyroid hormone-like peptide associated with humoral hypercalcemia of malignancy and parathyroid hormone bind to the same receptor on the plasma membrane of ROS 17/2.8 cells. J Biol Chem (1988) 263:8557–8560.

2. Fredriksson R, Lagerström MC, Lundin L-G, Schiöth HB. The G-protein-coupled receptors in the human genome form five main families. Phylogenetic analysis, paralogon groups, and fingerprints. Mol Pharmacol (2003) 63:1256–1272. doi:10.1124/mol.63.6.1256

3. Lagerström MC, Schiöth HB. Structural diversity of G protein-coupled receptors and significance for drug discovery. Nat Rev Drug Discov (2008) 7:339–357. doi:10.1038/nrd2518

4. Pioszak AA, Xu HE. Molecular recognition of parathyroid hormone by its G protein-coupled receptor. Proc Natl Acad Sci U S A (2008) 105:5034–5039. doi:10.1073/pnas.0801027105

5. Grauschopf U, Lilie H, Honold K, Wozny M, Reusch D, Esswein A, Schäfer W, Rücknagel KP, Rudolph R. The N-terminal fragment of human parathyroid hormone receptor 1 constitutes a hormone binding domain and reveals a distinct disulfide pattern. Biochemistry (2000) 39:8878–8887.

6. Karpf DB, Arnaud CD, Bambino T, Duffy D, King KL, Winer J, Nissenson RA. Structural properties of the renal parathyroid hormone receptor: hydrodynamic analysis and protease sensitivity. Endocrinology (1988) 123:2611–2620.

7. Ehrenmann J, Schöppe J, Klenk C, Rappas M, Kummer L, Doré AS, Plückthun A. High-resolution crystal structure of parathyroid hormone 1 receptor in complex with a peptide agonist. Nat Struct Mol Biol (2018) 25:1086–1092. doi:10.1038/s41594-018-0151-4

8. Castro M, Nikolaev VO, Palm D, Lohse MJ, Vilardaga J-P. Turn-on switch in parathyroid hormone receptor by a two-step parathyroid hormone binding mechanism. Proc Natl Acad Sci U S A (2005) 102:16084–16089. doi:10.1073/pnas.0503942102

9. Gensure RC, Gardella TJ, Jüppner H. Parathyroid hormone and parathyroid hormone-related peptide, and their receptors. Biochem Biophys Res Commun (2005) 328:666–678. doi:10.1016/j.bbrc.2004.11.069

10. Friedman PA. PTH revisited. Kidney Int Suppl (2004) S13–S19. doi:10.1111/j.1523-1755.2004.09103.x

11. Jüppner H, Abou-Samra AB, Freeman M, Kong XF, Schipani E, Richards J, Kolakowski LF, Hock J, Potts JT, Kronenberg HM. A G protein-linked receptor for parathyroid hormone and parathyroid hormone-related peptide. Science (1991) 254:1024–1026.

12. Vilardaga J-P, Romero G, Friedman PA, Gardella TJ. Molecular basis of parathyroid hormone receptor signaling and trafficking: a family B GPCR paradigm. Cell Mol Life Sci (2011) 68:1–13. doi:10.1007/s00018-010-0465-9

13. Cole JA. Parathyroid hormone activates mitogen-activated protein kinase in opossum kidney cells. Endocrinology (1999) 140:5771–5779. doi:10.1210/endo.140.12.7173

14. Mahon MJ, Bonacci TM, Divieti P, Smrcka AV. A docking site for G protein βγ subunits on the parathyroid hormone 1 receptor supports signaling through multiple pathways. Mol Endocrinol (2006) 20:136–146. doi:10.1210/me.2005-0169

15. Singh ATK, Gilchrist A, Voyno-Yasenetskaya T, Radeff-Huang JM, Stern PH. G alpha12/G alpha13 subunits of heterotrimeric G proteins mediate parathyroid hormone activation of phospholipase D in UMR-106 osteoblastic cells. Endocrinology (2005) 146:2171–2175. doi:10.1210/en.2004-1283

16. Malecz N, Bambino T, Bencsik M, Nissenson RA. Identification of phosphorylation sites in the G protein-coupled receptor for parathyroid hormone. Receptor phosphorylation is not required for agonist-induced internalization. Mol Endocrinol (1998) 12:1846–1856.

17. Gesty-Palmer D, Chen M, Reiter E, Ahn S, Nelson CD, Wang S, Eckhardt AE, Cowan CL, Spurney RF, Luttrell LM, et al. Distinct beta-arrestin- and G protein-dependent pathways for parathyroid hormone receptor-stimulated ERK1/2 activation. J Biol Chem (2006) 281:10856–10864. doi:10.1074/jbc.M513380200

18. Schwindinger WF, Fredericks J, Watkins L, Robinson H, Bathon JM, Pines M, Suva LJ, Levine MA. Coupling of the PTH/PTHrP receptor to multiple G-proteins. Direct demonstration of receptor activation of Gs, Gq/11, and Gi(1) by [alpha-32P]GTP-gamma-azidoanilide photoaffinity labeling. Endocrine (1998) 8:201–209. doi:10.1385/ENDO:8:2:201

19. Jilka RL. Molecular and cellular mechanisms of the anabolic effect of intermittent PTH. Bone (2007) 40:1434–1446. doi:10.1016/j.bone.2007.03.017

20. Lee M, Partridge NC. Parathyroid hormone signaling in bone and kidney. Curr Opin Nephrol Hypertens (2009) 18:298–302. doi:10.1097/MNH.0b013e32832c2264

21. Whitfield JF, Morley P, Willick GE, Ross V, Barbier JR, Isaacs RJ, Ohannessian-Barry L. Stimulation of the growth of femoral trabecular bone in ovariectomized rats by the novel parathyroid hormone fragment, hPTH-(1–31)NH2 (Ostabolin). Calcif Tissue Int (1996) 58:81–87.

22. Yang D, Singh R, Divieti P, Guo J, Bouxsein ML, Bringhurst FR. Contributions of parathyroid hormone (PTH)/PTH-related peptide receptor signaling pathways to the anabolic effect of PTH on bone. Bone (2007) 40:1453–1461. doi:10.1016/j.bone.2007.02.001

23. Hilliker S, Wergedal JE, Gruber HE, Bettica P, Baylink DJ. Truncation of the amino terminus of PTH alters its anabolic activity on bone in vivo. Bone (1996) 19:469–477.

24. Gesty-Palmer D, Flannery P, Yuan L, Corsino L, Spurney R, Lefkowitz RJ, Luttrell LM. A beta-arrestin-biased agonist of the parathyroid hormone receptor (PTH1R) promotes bone formation independent of G protein activation. Sci Transl Med (2009) 1:1ra1. doi:10.1126/scitranslmed.3000071

25. Potts JT. Parathyroid hormone: past and present. J Endocrinol (2005) 187:311–325. doi:10.1677/joe.1.06057

26. Klenk C, Schulz S, Calebiro D, Lohse MJ. Agonist-regulated cleavage of the extracellular domain of parathyroid hormone receptor type 1. J Biol Chem (2010) 285:8665–8674. doi:10.1074/jbc.M109.058685

27. Lupp A, Klenk C, Röcken C, Evert M, Mawrin C, Schulz S. Immunohistochemical identification of the PTHR1 parathyroid hormone receptor in normal and neoplastic human tissues. Eur J Endocrinol (2010) 162:979–986. doi:10.1530/EJE-09-0821

28. Klenk C, Vetter T, Zürn A, Vilardaga J-P, Friedman PA, Wang B, Lohse MJ. Formation of a ternary complex among NHERF1, beta-arrestin, and parathyroid hormone receptor. J Biol Chem (2010) 285:30355–30362. doi:10.1074/jbc.M110.114900

29. Emami-Nemini A, Gohla A, Urlaub H, Lohse MJ, Klenk C. The guanine nucleotide exchange factor Vav2 is a negative regulator of parathyroid hormone receptor/Gq signaling. Mol Pharmacol (2012) 82:217–225. doi:10.1124/mol.112.078824

30. Kumar S, Ratnikov BI, Kazanov MD, Smith JW, Cieplak P. CleavPredict: A Platform for Reasoning about Matrix Metalloproteinases Proteolytic Events. PLoS ONE (2015) 10:e0127877. doi:10.1371/journal.pone.0127877

31. Zhao L-H, Ma S, Sutkeviciute I, Shen D-D, Zhou XE, de Waal PW, Li C-Y, Kang Y, Clark LJ, Jean-Alphonse FG, et al. Structure and dynamics of the active human parathyroid hormone receptor-1. Science (2019) 364:148–153. doi:10.1126/science.aav7942

32. Hakalahti AE, Vierimaa MM, Lilja MK, Kumpula E-P, Tuusa JT, Petäjä-Repo UE. Human beta1-Adrenergic Receptor Is Subject to Constitutive and Regulated N-terminal Cleavage. J Biol Chem (2010) 285:28850–28861. doi:10.1074/jbc.M110.149989

33. Kozuka M, Ito T, Hirose S, Lodhi KM, Hagiwara H. Purification and characterization of bovine lung endothelin receptor. J Biol Chem (1991) 266:16892–16896.

34. Grantcharova E, Furkert J, Reusch HP, Krell H-W, Papsdorf G, Beyermann M, Schulein R, Rosenthal W, Oksche A. The extracellular N terminus of the endothelin B (ETB) receptor is cleaved by a metalloprotease in an agonist-dependent process. J Biol Chem (2002) 277:43933–43941. doi:10.1074/jbc.M208407200

35. Couet J, Sar S, Jolivet A, Hai MT, Milgrom E, Misrahi M. Shedding of human thyrotropin receptor ectodomain. Involvement of a matrix metalloprotease. J Biol Chem (1996) 271:4545–4552.

36. Kaczur V, Puskas LG, Nagy ZU, Miled N, Rebai A, Juhasz F, Kupihar Z, Zvara A, Hackler L, Farid NR. Cleavage of the human thyrotropin receptor by ADAM10 is regulated by thyrotropin. J Mol Recognit (2007) 20:392–404. doi:10.1002/jmr.851

37. Boire A, Covic L, Agarwal A, Jacques S, Sherifi S, Kuliopulos A. PAR1 is a matrix metalloprotease-1 receptor that promotes invasion and tumorigenesis of breast cancer cells. Cell (2005) 120:303–313. doi:10.1016/j.cell.2004.12.018

38. Ludeman MJ, Zheng YW, Ishii K, Coughlin SR. Regulated shedding of PAR1 N-terminal exodomain from endothelial cells. J Biol Chem (2004) 279:18592–18599. doi:10.1074/jbc.M310836200

39. Vallon M, Essler M. Proteolytically processed soluble tumor endothelial marker (TEM) 5 mediates endothelial cell survival during angiogenesis by linking integrin alpha(v)beta3 to glycosaminoglycans. J Biol Chem (2006) 281:34179–34188. doi:10.1074/jbc.M605291200

40. Mattila SO, Tuhkanen HE, Lackman JJ, Konzack A, Morató X, Argerich J, Saftig P, Ciruela F, Petäjä-Repo UE. GPR37 is processed in the N-terminal ectodomain by ADAM10 and furin. FASEB J (2021) 35:e21654. doi:10.1096/fj.202002385RR

41. Mattila SO, Tuusa JT, Petäjä-Repo UE. The Parkinson’s-disease-associated receptor GPR37 undergoes metalloproteinase-mediated N-terminal cleavage and ectodomain shedding. J Cell Sci (2016) 129:1366–1377. doi:10.1242/jcs.176115

42. Luttrell LM, Maudsley S, Bohn LM. Fulfilling the Promise of “Biased” G Protein-Coupled Receptor Agonism. Mol Pharmacol (2015) 88:579–588. doi:10.1124/mol.115.099630

43. Kenakin TP. New concepts in pharmacological efficacy at 7TM receptors: IUPHAR review 2. Br J Pharmacol (2013) 168:554–575. doi:10.1111/j.1476-5381.2012.02223.x

44. Onaran HO, Rajagopal S, Costa T. What is biased efficacy? Defining the relationship between intrinsic efficacy and free energy coupling. Trends Pharmacol Sci (2014) 35:639–647. doi:10.1016/j.tips.2014.09.010

45. Bisello A, Chorev M, Rosenblatt M, Monticelli L, Mierke DF, Ferrari SL. Selective ligand-induced stabilization of active and desensitized parathyroid hormone type 1 receptor conformations. J Biol Chem (2002) 277:38524–38530. doi:10.1074/jbc.M202544200

46. Takasu H, Gardella TJ, Luck MD, Potts JT, Bringhurst FR. Amino-terminal modifications of human parathyroid hormone (PTH) selectively alter phospholipase C signaling via the type 1 PTH receptor: implications for design of signal-specific PTH ligands. Biochemistry (1999) 38:13453–13460.

47. Jouishomme H, Whitfield JF, Chakravarthy B, Durkin JP, Gagnon L, Isaacs RJ, Maclean S, Neugebauer W, Willick G, Rixon RH. The protein kinase-C activation domain of the parathyroid hormone. Endocrinology (1992) 130:53–60. doi:10.1210/endo.130.1.1727720

48. Azarani A, Goltzman D, Orlowski J. Structurally diverse N-terminal peptides of parathyroid hormone (PTH) and PTH-related peptide (PTHRP) inhibit the Na+/H+ exchanger NHE3 isoform by binding to the PTH/PTHRP receptor type I and activating distinct signaling pathways. J Biol Chem (1996) 271:14931–14936.

49. Wheatley M, Wootten D, Conner MT, Simms J, Kendrick R, Logan RT, Poyner DR, Barwell J. Lifting the lid on GPCRs: the role of extracellular loops. Br J Pharmacol (2012) 165:1688–1703. doi:10.1111/j.1476-5381.2011.01629.x

50. White AD, Fang F, Jean-Alphonse FG, Clark LJ, An H-J, Liu H, Zhao Y, Reynolds SL, Lee S, Xiao K, et al. Ca2+ allostery in PTH-receptor signaling. Proc Natl Acad Sci U S A (2019) 116:3294–3299. doi:10.1073/pnas.1814670116

51. Li M, Li M, Guo J. Molecular Mechanism of Ca2+ in the Allosteric Regulation of Human Parathyroid Hormone Receptor-1. J Chem Inf Model (2021) doi:10.1021/acs.jcim.1c00471

52. Mitra N, Liu Y, Liu J, Serebryany E, Mooney V, Devree BT, Sunahara RK, Yan ECY. Calcium-dependent ligand binding and G-protein signaling of family B GPCR parathyroid hormone 1 receptor purified in nanodiscs. ACS Chem Biol (2013) 8:617–625. doi:10.1021/cb300466n

53. Koth CM, Murray JM, Mukund S, Madjidi A, Minn A, Clarke HJ, Wong T, Chiang V, Luis E, Estevez A, et al. Molecular basis for negative regulation of the glucagon receptor. Proc Natl Acad Sci U S A (2012) 109:14393–14398. doi:10.1073/pnas.1206734109

54. Liang Y-L, Khoshouei M, Deganutti G, Glukhova A, Koole C, Peat TS, Radjainia M, Plitzko JM, Baumeister W, Miller LJ, et al. Cryo-EM structure of the active, Gs-protein complexed, human CGRP receptor. Nature (2018) 561:492–497. doi:10.1038/s41586-018-0535-y

55. Dal Maso E, Glukhova A, Zhu Y, García-Nafría J, Tate CG, Atanasio S, Reynolds CA, Ramírez-Aportela E, Carazo J-M, Hick CA, et al. The Molecular Control of Calcitonin Receptor Signaling. ACS Pharmacology & Translational Science (2019) 2:31–51. doi:10.1021/acsptsci.8b00056

56. Liang Y-L, Khoshouei M, Glukhova A, Furness SGB, Zhao P, Clydesdale L, Koole C, Truong TT, Thal DM, Lei S, et al. Phase-plate cryo-EM structure of a biased agonist-bound human GLP-1 receptor–Gs complex. Nature (2018) 68:954. doi:10.1002/jcc.20084

57. Lei S, Clydesdale L, Dai A, Cai X, Feng Y, Yang D, Liang Y-L, Koole C, Zhao P, Coudrat T, et al. Two distinct domains of the glucagon-like peptide-1 receptor control peptide-mediated biased agonism. J Biol Chem (2018) 293:9370–9387. doi:10.1074/jbc.RA118.003278

58. Sarkar K, Joedicke L, Westwood M, Burnley R, Wright M, McMillan D, Byrne B. Modulation of PTH1R signaling by an ECD binding antibody results in inhibition of β-arrestin 2 coupling. Sci Rep (2019) 9:14432. doi:10.1038/s41598-019-51016-z

59. Clark LJ, Clark LJ, Krieger J, Krieger J, White AD, White AD, Bondarenko V, Bondarenko V, Lei S, Lei S, et al. Allosteric interactions in the parathyroid hormone GPCR–arrestin complex formation. Nat Chem Biol (2020) 42:946. doi:10.1007/s13361-011-0261-2

60. Christopoulos A, Christopoulos G, Morfis M, Udawela M, Laburthe M, Couvineau A, Kuwasako K, Tilakaratne N, Sexton PM. Novel receptor partners and function of receptor activity-modifying proteins. J Biol Chem (2003) 278:3293–3297. doi:10.1074/jbc.C200629200

61. Nemec K, Schihada H, Kleinau G, Zabel U, Grushevskyi EO, Scheerer P, Lohse MJ, Maiellaro I. Functional modulation of PTH1R activation and signalling by RAMP2. bioRxiv (2021) 2021.12.08.471790. doi:10.1101/2021.12.08.471790

62. Schechter I, Berger A. On the size of the active site in proteases. I. Papain. Biochem Biophys Res Commun (1967) 27:157–162. doi:10.1016/j.bbrc.2012.08.015

